# Dysglycemia is associated with *Mycobacterium tuberculosis* lineages in tuberculosis patients of North Lima - Peru

**DOI:** 10.1101/2020.11.18.388124

**Authors:** Kattya Lopez, María B. Arriaga, Juan G. Aliaga, Nadia N. Barreda, Oswaldo M. Sanabria, Chuan-Chin Huang, Zibiao Zhang, Ruth García-de-la-Guarda, Leonid Lecca, Anna Cristina Calçada Carvalho, Afrânio L. Kritski, Roger I. Calderon

## Abstract

This study was performed to investigate the role of dysglycemia on the genetic diversity of *Mycobacterium tuberculosis* (MTB) among pulmonary tuberculosis (TB) patients to build scientific evidence about the possible mechanisms of TB transmission. MTB isolates obtained of patients affected by pulmonary tuberculosis from health care facilities of North Lima - Peru, were analyzed using whole genome sequencing and 24-locus mycobacterial interspersed repetitive-unit-variable-number tandem repeats (MIRU-VNTR). Subsequently, clinical and epidemiological characteristics were associated with clustering, lineages and comorbid conditions. The analysis carried out 112 pulmonary TB patients from various health centers in North Lima, 17 (15%) had diabetes mellitus (DM) and 33 (29%) had pre-diabetes (PDM). Latin American-Mediterranean, Haarlem and Beijing were the most frequent MTB lineages found in those patients. Previous TB (adjusted odds ratio [aOR]=3.65; 95%CI: 1.32-17.81), age (aOR=1.12; 95%CI: 1.03-1.45) and Beijing lineage (aOR=3.53; 95%CI: 1.08-13.2) were associated with TB-DM comorbidity. Alcoholism (aOR=2.92; 95%CI: 1.10-8.28), age (aOR=1.05; 95%CI: 1.03-1.12) and Haarlem lineage (aOR=2.54; 95%CI: 1.04-6.51) were associated with TB-PDM comorbidity. Beijing and Haarlem lineages were independently associated with TB-DM and TB-PDM comorbidities, respectively. Although these findings may be surprising, we must be cautious to suggest that dysglycemia could be associated with a highly clustering and predisposition of MTB lineages related to a serious impact on the severity of TB disease, which requires further research.

## Introduction

Since recent years, dysglycemic conditions; known as diabetes mellitus (DM) and pre-diabetes (PDM), have been frequently reported in countries where tuberculosis (TB) is highly endemic [1–4]. In agreement with the World Health Organization, both TB and DM were two of the top ten causes of death worldwide in 2017 [1]. A strongly association between DM and worse TB treatment outcomes was widely documented [5, 6]. However, prediabetes (PDM), a state of increased risk of progression of diabetes, just also reported more likely to have the infective form of TB [7]. The comorbidity was not an important public health issue in low- and middle-income countries but the relevance of this situation has progressively changed due to urbanization and lifestyles changes [8]. Research in this regard is still very limited in Peru, which is why we are joining through different efforts to learn more about comorbidity [9] to try know if could impact in the risk of TB transmission and simultaneously explain the clinical burden of tuberculosis in the country.

In Peru, the Ministry of Health reported a prevalence of 6% of DM in TB patients in 2017 and more than 50% of these patients were concentrated in Lima [10]. Although information on comorbidity across different areas of Lima is still scarce, two recent studies reported a marked difference: 5.8% cases of TB-DM and 13% cases of TB-PDM [11], as well as, 14% cases of TB-DM and more than 30% cases of TB-PDM [12]. In most Peruvian health care facilities, despite increased comorbidity, the systematic diagnosis of dysglycemia in TB patients is complicated due to the testing cost and access barriers [12] and both, patients and physicians, remain unaware of this condition and the possible consequences. It is widely considered that unawareness of the comorbid conditions in these patients leads to poor management of the disease, with worse treatment outcomes and triggers us the intention to explore different strategies to improve control of the burden and transmission of TB.

*Mycobacterium tuberculosis* (MTB) genotyping helps to understand the clonal propagation and geographical distribution of TB strains [13], identifying the transmission of isolates with characteristics related with worse outcomes [14, 15], as virulence or drug resistance. However, the relevance of immunosuppressive characteristics on the spread or transmission of TB has not been formally documented. The dizzying spread of some MTB lineages in different populations plays an unclear role in disease outcome and makes us speculate on the human factors that predispose this specific transmissibility [16]. Since dysglycemia affects TB disease presentation [17] and treatment outcomes [18], we hypothesize that dysglycemia could predispose to a specific lineage and to influence on the TB burden in Peru.

To understand this problem, we assess the relation between dysglycemic status and MTB lineages in isolates of pulmonary TB patients from several health care facilities from North Lima. This research will contribute to get an insight of factors involved in transmission and help to control the TB in Peru.

## Materials and methods

### Ethics statement

The study protocol was approved by the Institutional Committee of Ethics for Humans (Approval number: 158-22-16) of the Universidad Peruana Cayetano Heredia accredited by the *Instituto Nacional de Salud* of Peru and the *Dirección de Redes Integradas de Salud de Lima Norte, Peru* (North Lima - Peru), to enroll patients in health care facilities. Written informed consents were obtained from all patients. The study was undertaken in agree with the principles of the Helsinki Declaration and Peruvian regulations.

### Study Settings

This cross-sectional study was performed using information from a larger prospective cohort conducted in North Lima aimed to explain the dysglycemia prevalence in pulmonary TB patients and their household contacts [12].

### Study population

For this study, TB patients enrolled were ≥18 years of age, diagnosed between February and November 2017 at 18 public health centers in Comas and Carabayllo, districts of North Lima. All health care facilities provided diagnosis and treatment supported by the local TB Program following guidelines of the Peruvian Ministry of Health [19]. Also, the enrolling process included patients who have not received anti-TB treatment or had started in no more than 5 days prior.

### Laboratory and field procedures

Microbiological procedures for TB diagnosis and drug susceptibility testing were performed at the Socios En Salud (SES) Laboratory following conventional recommendations as described [12, 19]. MTB isolates from culture processing were stored at −80°C until genotyping analysis.

DM was determined by an endocrinologist in agreement with American Diabetes Association (ADA) guidelines as fasting plasma glucose (FPG) ≥126 mg/dL, glycated hemoglobin (HbA1c) ≥6.5% and 2-h glucose ≥200 mg/dL of oral glucose tolerance test (OGTT). Also, PDM was determined in agreement with ADA guidelines as FPG 100 to 125 mg/dL, HbA1c 5.7 to 6.4% or 2-h glucose OGTT of 140 to 199 mg/dL [12, 20]. Additionally, SES laboratory undergoes an annual external quality control through panels from College of American Pathologists (Northfield, Illinois) and BC50 Clinical Chemistry Program of BioRad for Biochemistry and Hematology testing [12, 17].

### Clinical data

Epidemiological, anthropometric and clinical information were retrieved from interviews and medical records [12]. Variables such as age, sex, weight, height, body mass index (BMI), previous TB, Bacillus Calmette–Guerin (BCG) vaccine scar, TB symptoms, smear microscopy results, drug susceptibility test results, DM symptoms, hypoglycemic drugs and comorbidities were included. Chest radiographs were examined by the study staff graded the lung injury based on the number of cavitations and presence of fibrous tracts, alveolar infiltrate, pleural effusion or miliary dissemination, following the Peruvian guidelines [17, 19].

### Genotype analysis

MTB lineage assignment was based on Mycobacterial Interspersed Repetitive-Unit–Variable-Number Tandem-Repeat (MIRU-VNTR) using the reference database [21] and whole genome sequencing (WGS) [22]. The genomic sequences were mapped with the Burrows-Wheeler Aligner algorithm with maximal exact match seeds [23] against the MTB reference genome H37Rv. To identify single nucleotide polymorphisms, sequences were aligned using SAMtools [24] and Pilon [25]. Finally, MTB lineages were assigned based on WGS using SNP barcodes [22] as complement to MIRU-VNTR analysis.

The assignment of MTB lineages in our study population were determined by both 24-loci-MIRU-VNTR and WGS methods (Table 2) in 86 MTB isolates. Due in 26 isolates was not possible to have WGS profiles, were decide the assignment only using MIRU-VNTR method. This will be appropriate because this method offers adequate resolution and continues to be a genotyping method widely used in low and middle income areas [26]. Discordant MTB lineages assignments (n=16) were solved taking the information from WGS due its more robust and discriminatory classification [27].

### Outcomes

The primary outcome was the relation between MTB lineage and dysglycemic status in TB patients.

### Statistical analysis

Continuous variables were presented as median and interquartile range (IQR) values and were compared using the Mann-Whitney U test (between two groups) or the Kruskal-Wallis test with Dunn’s multiple comparisons (between >2 groups). Categorical variables were presented by frequency and compared using the Fisher’s exact test (2×2 comparisons) or Pearson’s chi square test.

We also performed Kappa (k) statistic to determined agreement between 24-MIRU-VNTR and WGS using five levels of agreement: <0.20 (poor), 0.21–0.40 (weak), 0.41–0.60 (moderate), 0.61–0.80 (good), and 0.81–1.00 (very good) [12].

Variables with univariate p-value ≤ 0.2 were used in the generalized linear mixed-effects model analysis, where the “health care facilities” variable was included as a repetitive measure to assess the odds ratios (OR) and 95% confidence intervals (CIs) of the associations between MTB lineage among TB-patients with DM and PDM. Two-sided p-value < 0.05 were considered statistically significant.

The hierarchical clustering was performed as additional analysis using the 24-loci-MIRU-VNTR and based on categorical distances as described [21]. Briefly, a genotypic cluster was defined as a group of two or more isolates from different patients with identical 24-MIRU-VNTR profiles. The lineage-clustering rate was calculated between the difference of the total number of strain-clustered cases in each MTB lineage and the number of clusters, divided by the total number of cases in the MTB lineage.

Statistical analysis was performed using SPSS version 24.0 (IBM statistics), Graphpad Prism 7.0 (GraphPad Software, San Diego, CA) and JMP 13.0 (SAS, Cary, NC, USA).

## Results

This study included 112 pulmonary TB patients, 17 (15%) were affected by DM, 33 (29%) by PDM. Both TB-DM and TB-PDM patients were significantly older than patients with only TB (average of 10.0 years older, p<0.01). Additionally, they had similar distribution in BMI, previous TB and BCG vaccination. TB-DM patients were more likely to have hypertension than non-DM patients (p<0.01). Furthermore, TB-DM and TB-PDM patients showed significantly more lung lesions (either cavitation, fibrous tracts, alveolar infiltrate, pleural effusion or miliary dissemination) (p<0.01) and significantly higher bacillary load than normoglycemic patients (p<0.001). On the other hand, there was no significant difference in TB symptoms such as fever>15 days, cough>15 days, blood in sputum and night sweats between patients with only TB and TB patients with dysglycemia (Table 1). However, weight loss was significantly more frequent among TB-DM patients compared to TB-PDM patients (p<0.03). Likewise, we found no statistically significant differences in the proportions of MDR-TB patients the study: (18% in DM-TB, 13% in TB-PDM and 10% in only TB) (Table 1). Other characteristics are shown in Supplementary Table 1.

**Table 1.** Baseline epidemiological and clinical characteristics of the study population.

| Characteristics<sup>++</sup> | Total<br>(N=112) | TB-DM<br>(n=17) | TB-PDM<br>(n=33) | Only TB<br>(n=62) | p-value* |
| --- | --- | --- | --- | --- | --- |
| ◊Age, years | 28.8<br>(23.1-67.7) | 46.4<br>(37.9-54.0) | 34.9<br>(25.2-48.7) | 25.7<br>(21.7-31.0) | <0.01 |
| Male | 69 (61.6) | 8 (47.1) | 22 (66.7) | 39 (62.9) | 0.39 |
| ◊BMI, kg/m<sup>2</sup> | 22.7<br>(25.1-27.9) | 22.3<br>(21.6-25.7) | 22.9<br>(20.5-25) | 22.6<br>(20.3-25.1) | 0.92 |
| Previous TB | 18 (16.1) | 5 (29.4) | 6 (18.2) | 7 (11.3) | 0.07 |
| BCG vaccination scar | 104 (93.7) | 15 (88.2) | 30 (93.8) | 59 (95.2) | 0.33 |
| Fever>15 days | 53 (47.3) | 5 (29.4) | 18 (54.5) | 30 (48.4) | 0.34 |
| Cough > 15 days | 103 (92.0) | 16 (94.1) | 30 (90.9) | 57 (91.9) | 0.86 |
| Weight loss | 81 (72.3) | 14 (82.4) | 28 (84.8) | 39 (62.9) | **0.03** |
| Night sweats | 69 (61.6) | 11 (64.7) | 19 (57.6) | 39 (62.9) | 0.94 |
| +◊Lung injury | 4 (2-5) | 7 (5-8) | 4.5 (3-5) | 2 (1-4) | <0.01 |
| °Smear result |  |  |  |  | **0.001** |
| Negative | 40 (35.7) | 3 (17.6) | 7 (21.2) | 30 (48.4) |  |
| Scanty or 1+ | 31 (27.7) | 4 (23.5) | 10 (30.3) | 17 (27.4) |  |
| 2+ | 14 (12.5) | 3 (17.6) | 6 (18.2) | 5 (8.1) |  |
| 3+ | 27 (24.1) | 7 (41.2) | 10 (30.3) | 10 (16.1) |  |
| Hypertension | 6 (5.4) | 4 (23.5) | 2 (6.3) | 0 (0) | <0.01 |

Data represent no. (%). TB=pulmonary tuberculosis, DM=diabetes mellitus, TB-DM=pulmonary tuberculosis and diabetes mellitus comorbidity, TB-PDM=pulmonary tuberculosis and prediabetes comorbidity at screening, BMI=body mass index.
+It was defined as the total number of lesions in the lung either cavitation, fibrous tracts, alveolar infiltrate, pleural effusion or miliary dissemination.
°All smear MTB positive results were stratified as scanty (up to 9 AFB), 1+ (10–99 AFB observed in 100 fields), 2+ (1 a 10 AFB in 50 fields) and 3+ (more than 10 AFB in 20 fields). AFB= Acid-fast bacillus
◊Continuous values are expressed as medians with interquartile intervals.
\*In black indicate p<0.05.
++Other characteristics are shown in S1 Table: symptoms of TB (dyspnea, blood in sputum, night sweats), multidrug resistance, DM symptoms (polydipsia, polyuria, fatigue, delayed wound healing), dysglycemia control drug use (metformin, insulin, other drugs), other comorbidities (alcoholism, smoking).

The assignment of MTB lineages by using both molecular methods showed a good agreement (Cohen’s kappa=0.7) among 86 isolates (Supplementary Table 2) and the distribution between the dysglycemic status in TB patients is shown in Table 2. Latin American-Mediterranean (LAM) (35%), Haarlem (24%) and Beijing (16%) were the most frequent MTB lineages in all TB patients in our study. Beijing was the most frequent MTB lineage in TB-DM patients (35%), whereas Haarlem (39%) and LAM (38%) were the most frequent MTB lineages among patients affected by TB-PDM and only TB, respectively. In the multivariate analysis (Table 3), we showed that Beijing lineage (OR=3.53; 95%CI: 1.08-13.2), previous TB (OR=3.65; 95%CI: 1.32-17.81) and age in years (OR=1.12; 95%CI: 1.03-1.45) were independently associated with TB-DM. But in the Haarlem lineage (OR=2.54; 95%CI: 1.04-6.51), alcoholism (OR=2.92; 95%CI: 1.10-8.28) and age in years (OR=1.05; 95%CI: 1.03-1.12) were independently associated with TB-PDM.

**Table 2.** *M. tuberculosis* lineage distributions of the study population.

| Characteristics | Total<br>(N=112) | TB-DM<br>(n=17) | TB-PDM<br>(n=33) | Only TB<br>(n=62) | p-value** |
| --- | --- | --- | --- | --- | --- |
| MTB lineage |  |  |  |  | 0.12 |
| Beijing | 18 (16.1) | 6 (35.3) | 2 (6.1) | 10 (16.1) | **0.03** |
| LAM | 40 (35.7) | 5 (29.4) | 11 (33.3) | 24 (38.7) | 0.73 |
| Haarlem | 27 (24.1) | 1 (5.9) | 13 (39.4) | 13 (21.0) | **0.02** |
| S-clade | 10 (8.9) | 2 (11.8) | 4 (12.1) | 4 (6.5) | 0.59 |
| Other Euro-American | 9 (8.0) | 2 (11.8) | 2 (6.1) | 5 (8.1) | 0.38 |
| Others* | 8 (7.1) | 1 (5.9) | 1 (3.0) | 6 (9.7) | 0.48 |

Data represent no. (%). TB=tuberculosis, DM=diabetes mellitus, TB-DM=pulmonary tuberculosis and diabetes mellitus comorbidity, TB-PDM=pulmonary tuberculosis and prediabetes comorbidity at screening, MTB=Mycobacterium tuberculosis \*Included: H37Rv, X, Ghana, S, S-type \*\*Chi-square test

**Table 3.** Factors associated with TB patients affected with diabetes mellitus or prediabetes.

| ◊Factors | N | Unadjusted OR<br>(95% CI) | p-value* | Adjusted OR<br>(95% CI) | p-value* |
| --- | --- | --- | --- | --- | --- |
| **+TB-DM comorbidity** |  |  |  |  |  |
| Previous TB | 16 | 3.30 (0.89-12.09) | 0.07 | 3.65 (1.32-17.81) | **0.03** |
| Age, years | 17 | 1.14 (1.07-1.21) | **<0.01** | 1.12 (1.03-1.45) | **<0.01** |
| MTB Beijing lineage | 12 | 2.84 (0.85-9.45) | **0.03** | 3.53 (1.08-13.2) | **0.03** |
| **++TB-PDM comorbidity** |  |  |  |  |  |
| Alcoholism | 53 | 2.20 (0.90-5.40) | 0.12 | 2.92 (1.10-8.28) | **0.02** |
| Age, years | 33 | 1.07 (1.03-1.11) | **<0.01** | 1.05 (1.03-1.12) | **<0.01** |
| MTB Haarlem lineage | 24 | 2.45 (0.97-6.20) | 0.05 | 2.54 (1.04-6.51) | **0.02** |

◊Results from the generalized linear mixed-effects model that displayed univariate p-value ≤ 0.2. The variable ‘health care facilities’ was used as a repetitive measure. OR: odds ratio; CI=confidence interval; TB=pulmonary tuberculosis, DM=diabetes mellitus, TB-DM=pulmonary tuberculosis and diabetes mellitus comorbidity, TB-PDM=pulmonary tuberculosis and prediabetes comorbidity at screening. MTB: Mycobacterium tuberculosis +The modelling used n = 6 corresponding to TB-DM patients infected with the MTB Beijing lineage. ++The modelling used n=13 corresponding to TB-PDM patients infected with the MTB Haarlem lineage. \*In black indicate p<0.05.

Finally, isolates belonging to Beijing lineage exhibited a greater clustering (72% of clustering rate) and both LAM (36% of clustering rate), Haarlem (40% of clustering rate) and other lineages showed clusters of up to 6 strains.

## Discussion

This study represents the first description of the MTB lineages distribution in TB patients affected by dysglycemia in Lima-Peru, through 24-loci-MIRU-VNTR and WGS. Our findings showed that TB-DM and TB-PDM patients of several health care facilities from Lima Norte were more likely to be infected by Beijing and Haarlem, respectively. The hierarchical analysis displayed an overall clustering rate of 14% of all TB isolates and Beijing lineage showed a high clustering rate (72%), compared to other studies [28, 29]. However, the clustering rate of our TB patients without dysglycemia was 25%, similar result observed in a previous study [29]. Although several Peruvian studies carried out on isolates of resistant-TB isolates have shown an important predominance of the Haarlem and LAM lineages, with a trend to increase the prevalence of the Haarlem genotype in Lima in the last decade [30–34], no more recent data have been reported. Nevertheless, the generalized predominance of these lineages in Peru could be due to a recent transmission. The Beijing family in Peru has been found in increasing proportion in recent years [32, 35], possibly related to Asian immigration in Peru since the last century. The prevalence of the Beijing lineage of 16.1% observed in this study is very similar to that found previously [35], which together with a high clustering rate, confirms the trend of greater transmission, which requires a possible explanation From that information obtained, we can hypothesize about the existence of mechanisms that allow the transmission of MTB lineages among affected patients with the same comorbidity or that they can be more susceptible to exogenous infection from other patients. As LAM is the most prevalent MTB lineage in Lima [15, 35], possibly the concomitant condition of DM or PDM will lead immunologic changes that increase the patient susceptibility to infection by Beijing or Haarlem lineages, developing active TB [13, 14], even being the less prevalent lineage in North Lima.

The generalized linear mixed-effects model showed that MTB Beijing lineage, age and previous TB were associated with TB-DM comorbidity. Also, MTB Haarlem lineage, age and alcoholism were associated with TB-PDM comorbidity. In addition, given the high degree of clustering in our patients, we include an analysis of the North Lima health centers distribution in our patients as a variable to investigate whether sharing common areas could have an impact on such association but, its effect was observed. Therefore, recognizing that our findings may be surprising, we can indicate that dysglycemia can strongly explain the adverse treatment outcomes and that the disinterest of this condition in patients with TB puts the efficacy of health interventions in programs at risk. of TB control in at least North Lima, since they still have a high incidence of TB. However, the study team understands that these findings require further investigation.

Regardless of dysglycemia type, immunocompromised people (elderly and alcoholics, among others) are at the highest risk of developing TB with possible poor treatment outcomes [36]. However, it remains controversial whether age is a risk factor for TB-DM comorbidity or whether is a confounding variable [36]. All other risk factors such as alcoholism and previous TB, should be considered as one of the first screening strategies of patients with TB or DM to ensure appropriate outcomes. Nevertheless, our major concern was the distribution of MTB Beijing lineage among TB-DM patients because such lineage has gained relevance in TB dissemination due to its association with pathogenicity and multidrug resistant [14, 37].

This study used data based on MIRU-VNTR, WGS and molecular clustering to address recent transmission profiles, which is very useful for evaluating the effectiveness of TB control programs (44–46). The idea that genetically diverse strains display distinct transmission dynamics even within the same community [38], could support the hypothesis that there was a genetic predisposition of TB patients with dysglycemia to be infected by Beijing lineage; at least in North Lima. In our study, we integrated a conscious diagnosis of dysglycemia against the molecular diversity of MTB, to establish better relationships and risk factors associated with transmission and to allow a better understanding of the TB burden.

On the other hand, consistent with previous studies, our TB-DM and TB-PDM patients could be more likely to be older and hypertensive than normoglycemic TB patients [11], so medical caregivers should take both into account to ensure better outcomes [39]. In addition, our TB patients with or without dysglycemia showed no difference in TB symptoms, contrary to other studies [40]. Therefore, the detection of DM or PDM in TB patients becomes a challenge for health workers since there is not differential symptomatology among comorbid patients, which could lead to a delay in the onset of treatment.

Moreover, patients with dysglycemia showed higher bacillary burden compared to TB patients without dysglycemia, similar to others [36, 40, 41]. This supports our hypothesis that TB patients with altered glucose metabolism may be less able to eliminate MTB [36], being a transmission-related concern. In our study, TB patients with dysglycemia were more likely to present severe lung injury, [17] as well as, in other foreign studies [42]. Despite the fact that radiographic presentation of TB could be influenced by different factors such as duration of illness and host immune response in patients without DM and with DM [43], we should consider this report as valuable information to address new questions around TB severity and the role of transmission risk of more virulent MTB strains [17].

This study had several limitations: First, the study comprised two districts of North Lima, which could represent a possible selection bias on the MTB lineages distribution and clinical characteristics of Lima or Peruvian population. Second, the design and inclusion criteria (only positive cultures for MTB) had a substantial impact on the number of assessed patients, likewise, higher bacillary load in TB-DM patients compared to normoglycemic TB patients may be over-represented by the same reason. Third, the radiographic information was limited because the local TB strategies did not use standard scores to grade pulmonary injuries, as previously reported [44]. Finally, all MTB isolates were not analyzed by both molecular methods; so the remained analysis was made by use of MIRU-VNTR, which offers adequate resolution and continues to be a genotyping method widely used in low and middle-income areas [26].

## Conclusion

In summary, our findings give us an insight of MTB lineages distribution among TB patients affected by DM or PDM from some health care facilities of North Lima. Beijing lineage was predominant in TB-DM patients while Haarlem lineage was associated with TB-PDM and both families showed the greatest probability to cluster. The high rate of clustering indicates active transmission of *M. tuberculosis* among dysglycemic patients, associated with previous TB and other characteristics. Nevertheless, to confirm our hypothesis, further community-based studies, should aim to study the transmission of specific strains in the population affected or not with dysglycemia, as well as, studies for causal inference will be needed to test whether some MTB lineages lead to high blood sugar levels. Interventions to prevent the increase of TB-DM or TB-PDM patients should focus on risk factors associated with TB treatment outcomes.

## Acknowledgements

We thank the field workers, staff of health care facilities staff, lab staff of Socios En Salud Sucursal Peru and specially, each patient who supported us in favor of research in our country. In addition, thanks to Ildiko Van Rhijn and Sara Suliman, for their support in the development of this research work.

---

**Figure 1.**
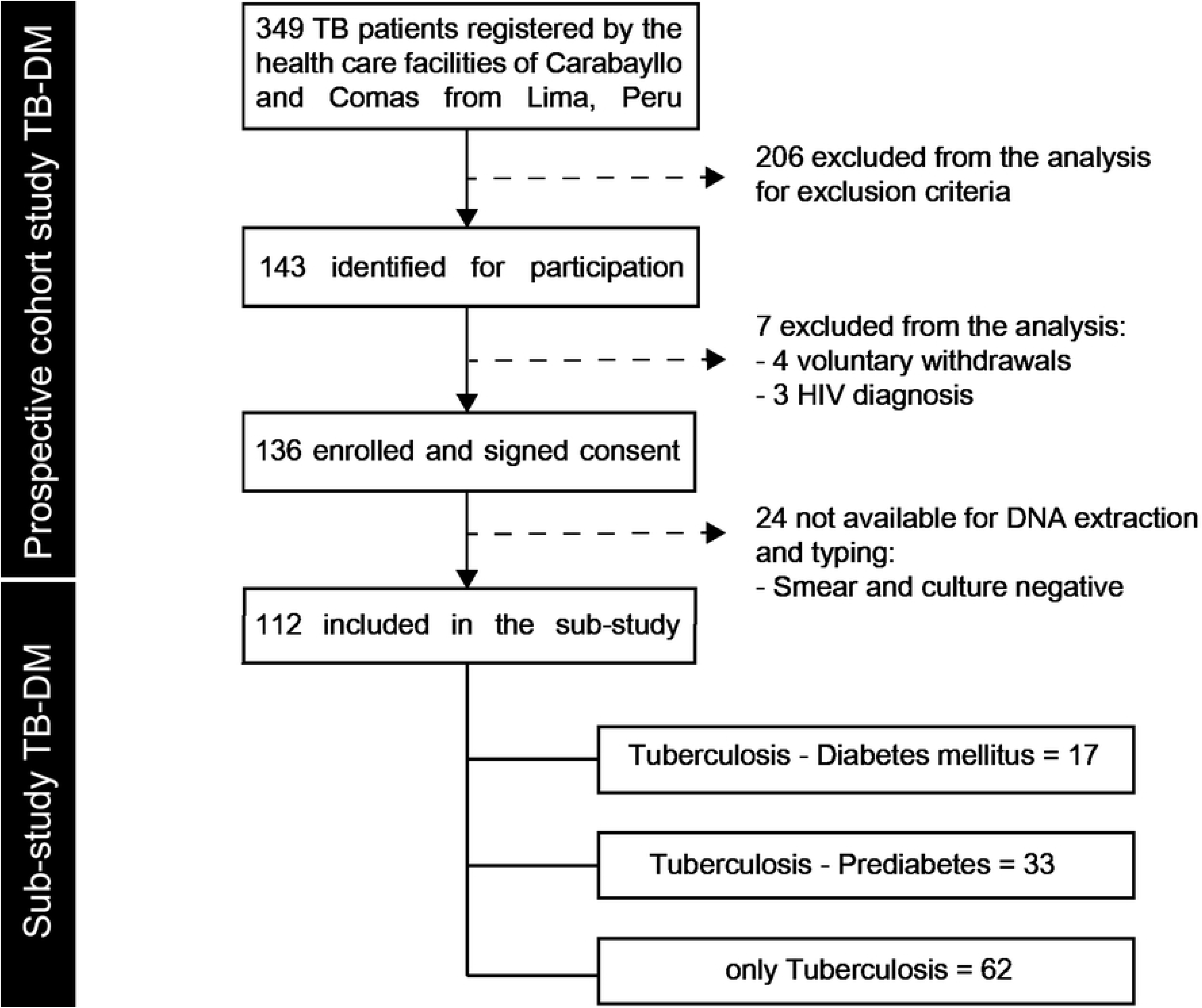
Flowchart of the study population. TB: pulmonary tuberculosis, DM: diabetes mellitus, PDM: prediabetes.

**Figure 2.**
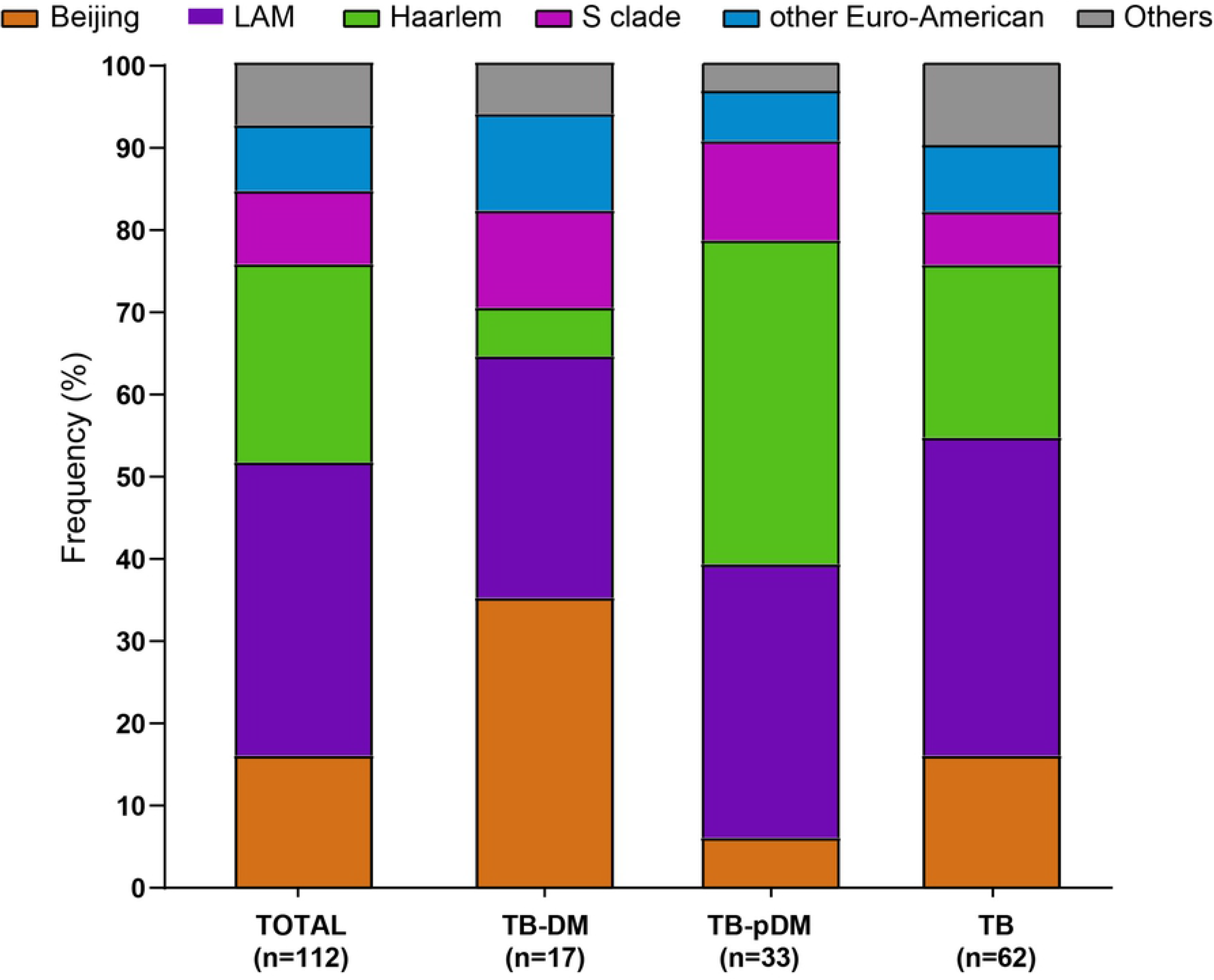
Frequency distribution of *M. tuberculosis* strain lineages among TB patients with diabetes, prediabetes and normal glucose levels (normoglycemic). Bar diagrams show the frequency in percentage of each lineage in each of these three categories. *M. tuberculosis* strain lineage assignments were based on 24-MIRU-VNTR analysis and whole genome sequencing (see method). The lineages H37Rv, X, Ghana, S, S-type were grouped under the label of “others” as the corresponding strains are very few in numbers.

**Figure 3.**
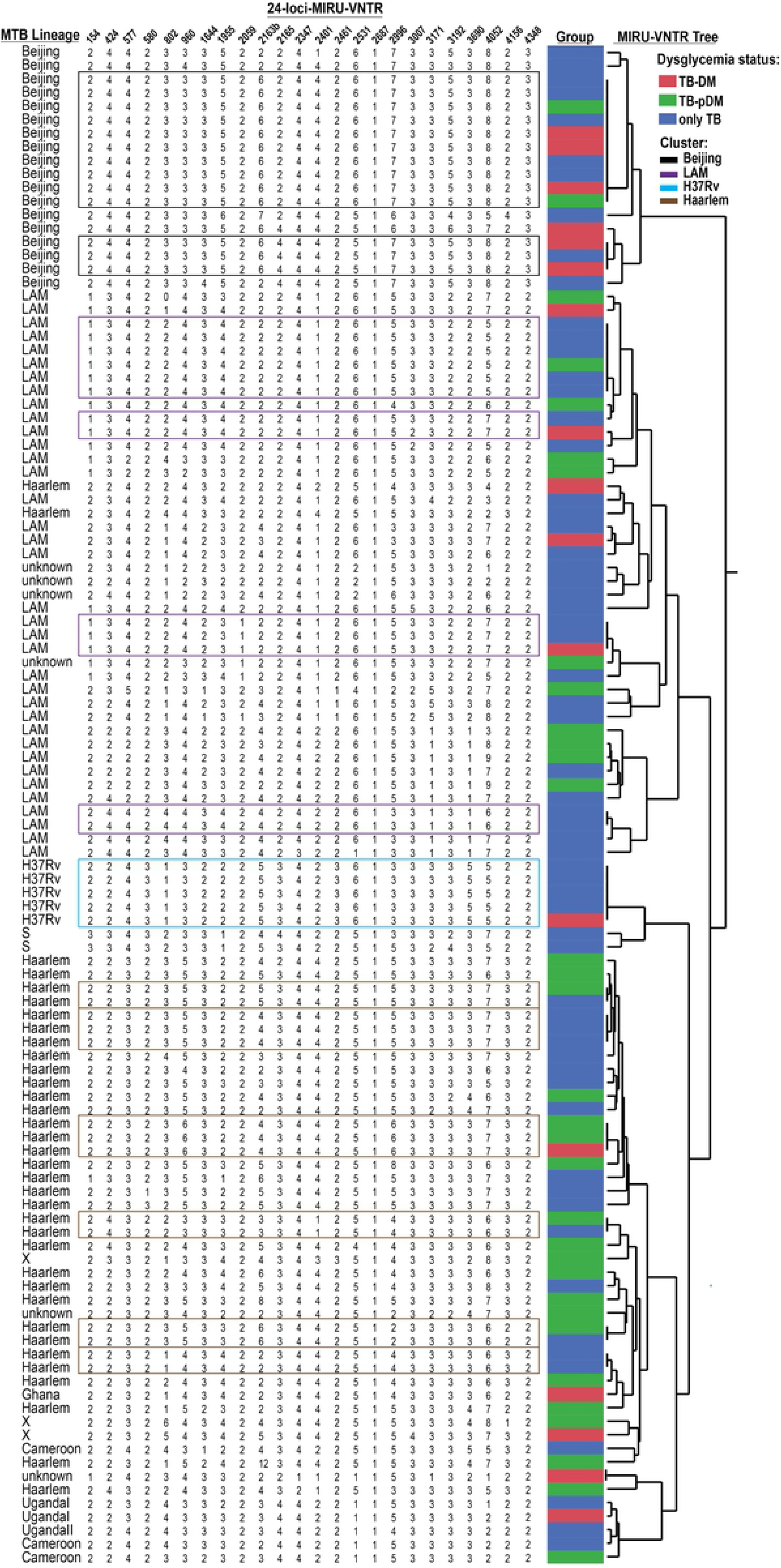
Cluster analysis of *M. tuberculosis* from clinical isolates of the study population. The MTB assignment and clustering analysis are based only on mycobacteria interspersed repetitive unit variable number tandem repeat analysis profiles among patients with tuberculosis and diabetes mellitus (TB-DM) (in red), patients with tuberculosis and prediabetes (TB-PDM) (in green) and patients with just TB alone (in blue). Some MTB isolates have shared clades and may be due to the presence of amplicons not determined by MIRU-VNTR and the most likely value assignment by JMP 13.0 software (SAS, Cary, NC, USA) has allowed limited discrimination; so WGS was necessary to confirm MTB lineages.

## Supporting information

**Supplementary Table 1. Baseline epidemiological and clinical characteristics of the study population.**

**Supplementary Table 2. Agreement analysis between 24-MIRU-VNTR and whole genome sequencing among 86 *M. tuberculosis* strains.**

